## Supplemental data for "Surface chemistry-mediated modulation of adsorbed albumin folding state specifies nanocarrier clearance by distinct macrophage subsets"

### Supplementary Tables.

**Supplementary Table 1. | Representative PEG-*b*-PPS polymers used in this study.**

| Block copolymer (BCP) | $f_{PEG}$[1] | MW (g/mol) |
|---|---|---|
| Phos-$PEG_{23}$-*b*-$PPS_{39}$-Bn | 0.21 | 3,911 |
| HO-$PEG_{23}$-*b*-$PPS_{40}$-Bn | 0.24 | 4,023 |
| MeO-$PEG_{17}$-*b*-$PPS_{28}$-Bn | 0.26 | 2,982 |

[1]Hydrophilic weight fraction of polymer.

**Supplementary Table 2. | Summary of PEG-*b*-PPS polymersome characterization by small angle x-ray scattering using synchrotron radiation.**

| Nanocarrier | SAXS | | |
|---|---|---|---|
| | Total Radius[1] (nm) | Core Radius[1] (nm) | Shell Thickness[1] (nm) |
| Phos PS | 38.2 | 28.7 | 9.5 |
| OH PS | 55.1 | 45.8 | 9.3 |
| MeO PS | 49.9 | 41.2 | 8.7 |

[1]Radii values (total radius, $r_t$; core radius, $r_c$; shell thickness, $r_{st}$) obtained after fitting SAXS profiles using a spherical vesicle model. $X^2$ of 0.13, 0.39, and 0.29 were obtained for models fit to Phos PS, OH PS, and MeO PS, respectively.

**Supplementary Table 3. | Cellular definitions and markers.**

| Organ | Cell Type | Abbreviation | Markers[1] |
|---|---|---|---|
| Spleen | Splenic non-immune cells | CD45- | CD45- |
| | CD169+ macrophages | MΦ (CD169+) | CD45+, CD3-, CD19-, NK1.1-, Ly-6C-, CD169+ |
| | Macrophages | MΦ (F4/80+) | CD45+, CD3-, CD19-, NK1.1-, Ly-6C-, CD169-, CD11c-, F4/80+, CD11b+ |
| | Dendritic cells | DC | CD45+, CD3-, CD19-, NK1.1-, CD11b-, CD11c+ |
| | Patrolling monocytes | Mono (Ly-6C$^{lo}$) | CD45+, CD3-, CD19-, NK1.1-, CD11b+, Ly-6C$^{lo}$, SSC$^{lo}$ |
| | Inflammatory monocytes | Mono (Ly-6C$^{hi}$) | CD45+, CD3-, CD19-, NK1.1-, CD11b+, Ly-6C$^{int}$, SSC$^{hi}$ |
| | Neutrophils & Eosinophils | Neu & Eos (Ly-6C$^{int}$) | CD45+, CD3-, CD19-, NK1.1-, CD11b+, Ly-6C$^{hi}$, SSC$^{lo}$ |
| Liver | Liver non-immune cells | CD45- | CD45- |
| | Macrophages & Kupffer cells | MΦ/K | CD45+, CD3-, CD19-, NK1.1-, Ly-6G-, CD11c-, CD11b+, F4/80+ |
| | Dendritic cells | DCs | CD45+, CD3-, CD19-, NK1.1-, Ly-6G-, CD11c+, IA/IE+ |
| | Monocytes | Mono | CD45+, CD3-, CD19-, NK1.1-, Ly-6G-, CD11c-, CD11b+, F4/80- |
| | Neutrophils | Neu | CD45+, CD3-, CD19-, NK1.1-, Ly-6G+ |
| LN | Lymph node non-immune cells | CD45- | CD45- |
| | Medullary sinus macrophages | MSMΦ | CD45+, CD3-, CD19-, NK1.1-, CD11b+, F4/80+, CD169+ |
| | Medullary cord macrophages | MCMΦ | CD45+, CD3-, CD19-, NK1.1-, CD11b+, F4/80+, CD169- |
| | Subcapsular sinus macrophages | SSMΦ | CD45+, CD3-, CD19-, NK1.1-, CD11b+, F4/80-, CD169+ |
| | Dendritic cells | DCs | CD45+, CD3-, CD19-, NK1.1-, CD11c+ |
| Kidney | Kidney non-immune cells | CD45- | CD45- |
| | Macrophages | MΦ | CD45+, CD3-, CD19-, NK1.1-, Ly-6G-, F4/80+, CD11b+ |
| | Dendritic cells | DC | CD45+, CD3-, CD19-, NK1.1-, Ly-6G-, F4/80-, CD11c+ |
| | Granulocytes | G (Ly-6G$^{hi}$) | CD45+, CD3-, CD19-, NK1.1-, Ly-6G+ |

[1]Gates were set on the population of live single cells using FSC, SSC, and the zombie violet viability stain.

**Supplementary Table 4. | Cellular definitions and markers used to define immune cell subpopulations in murine lung tissue.**

| Cell Type | Abbreviation | Markers[1,2] |
|---|---|---|
| Lung non-immune cells | CD45- | CD45- |
| Alveolar macrophages | AMΦ | CD45+, Ly-6C$^{-/lo/int}$, IA/IE+ or SSC$^{int/hi}$, CD24-, CD64+, CD11b-, CD11c+ |
| Interstitial macrophages | iMΦ | CD45+, Ly-6C$^{-/lo/int}$, IA/IE+ or SSC$^{int/hi}$, CD24-, CD64+, CD11b+, CD11c- |
| Dendritic cells | DC | CD45+, Ly-6C$^{-/lo/int}$, IA/IE+ or SSC$^{hi}$, CD64-, CD24+, CD11b-, IA/IE+ |
| Resident monocytes | rMono | CD45+, Ly-6C$^{-/lo/int}$, IA/IE-, SSC$^{lo}$, CD64$^{int}$, CD11b$^{hi}$, Ly-6C-, CD11c+ |
| Inflammatory monocytes | Mono (Ly-6C$^{hi}$) | CD45+, Ly-6C$^{-/lo/int}$, IA/IE-, SSC$^{lo}$, CD64$^{int}$, CD11b$^{hi}$, Ly-6C$^{hi}$, CD11c- |
| Granulocytes | G* (Ly-6C$^{hi}$)[3] | CD45+, CD11b-, CD11c-, Ly-6C$^{hi}$, SSC$^{hi}$ |
| Eosinophils | Eos | CD45+, Ly-6C$^{-/lo/int}$, IA/IE+ or SSC$^{hi}$, CD64-, CD24+, CD11b+, IA/IE- |
| B cells | B cells | CD45+, CD11b-, CD11c-, Ly-6C$^{-/lo/int}$, IA/IE+, CD24+ |
| T cells | T cells | CD45+, CD11b-, CD11c-, Ly-6C$^{-/lo/int}$, IA/IE-, CD24- |
| Natural killer cells | NK | CD45+, Ly-6C$^{-/lo/int}$, IA/IE-, SSC$^{lo}$, CD64-, CD11b$^{int}$ |

[1]Gates were set on the population of live single cells using FSC, SSC, and the zombie violet viability stain.
[2]This gating strategy follows that reported by Yu et al., 2016, with minor changes.
[3]The asterisk (G*) indicates this granulocyte does not include the explicitly defined eosinophil subpopulation.

**Supplementary Table 5. | Two-way ANOVA assessment of input BSA concentration and PEG-*b*-PPS PS surface chemistry on the concentration of BSA adsorbed onto PS nanocarriers.**

| Source of variation | Percentage of total variation | *p* value | *p* value summary | Significant? (alpha = 0.05) | |
|---|---|---|---|---|---|
| Interaction | 4.254 | 0.0952 | ns | No | |
| Row Factor (input BSA concentration, in. mg/mL) | 73.1 | <0.0001 | **** | Yes | |
| Column Factor (PS surface chemistry) | 14.43 | 0.0001 | *** | Yes | |
| **ANOVA table** | **SS** | **DF** | **MS** | **F (DFn, DFd)** | ***p* value** |
| Interaction | 57.89 | 4 | 14.47 | F (4, 18) = 2.33 | *p*=0.0952 |
| Row Factor (input BSA concentration, in. mg/mL) | 994.8 | 2 | 497.4 | F (2, 18) = 80.09 | *p*<0.000 |
| Column Factor (PS surface chemistry) | 196.3 | 2 | 98.17 | F (2, 18) = 15.81 | *p*=0.0001 |
| Residual | 111.8 | 18 | 6.211 | | |

The analysis was performed on the adsorbed protein data presented in **Fig. 3** of the main text (n=3). The effect of two factors, the input BSA concentration used during pre-incubation ([BSA] = 0.1, 1.0, or 10.0 mg/mL) and the PS surface chemistry (Phos, OH, and MeO), on the adsorbed protein concentration was assessed.

**Supplementary Table 6. | Tukey's multiple comparison test performed on the BSA percent α-helicity determined by CD spectroscopy.**

| Comparison | Mean diff. | 95.00% CI of diff. | Significant? | Summary | Adjusted *p* value |
|---|---|---|---|---|---|
| BSA vs. Heat | 16.31 | 14.21 to 18.41 | Yes | **** | <0.0001 |
| BSA vs. Phos | -3.916 | -6.015 to -1.817 | Yes | *** | 0.0008 |
| BSA vs. OH | 21.35 | 19.25 to 23.45 | Yes | **** | <0.0001 |
| BSA vs. MeO | 14.5 | 12.4 to 16.6 | Yes | **** | <0.0001 |
| Heat vs. Phos | -20.23 | -22.33 to -18.13 | Yes | **** | <0.0001 |
| Heat vs. OH | 5.04 | 2.941 to 7.139 | Yes | **** | <0.0001 |
| Heat vs. MeO | -1.814 | -3.913 to 0.2846 | No | ns | 0.0994 |
| Phos vs. OH | 25.27 | 23.17 to 27.37 | Yes | **** | <0.0001 |
| Phos vs. MeO | 18.41 | 16.31 to 20.51 | Yes | **** | <0.0001 |
| OH vs. MeO | -6.854 | -8.953 to -4.756 | Yes | **** | <0.0001 |

### Supplementary Figures.

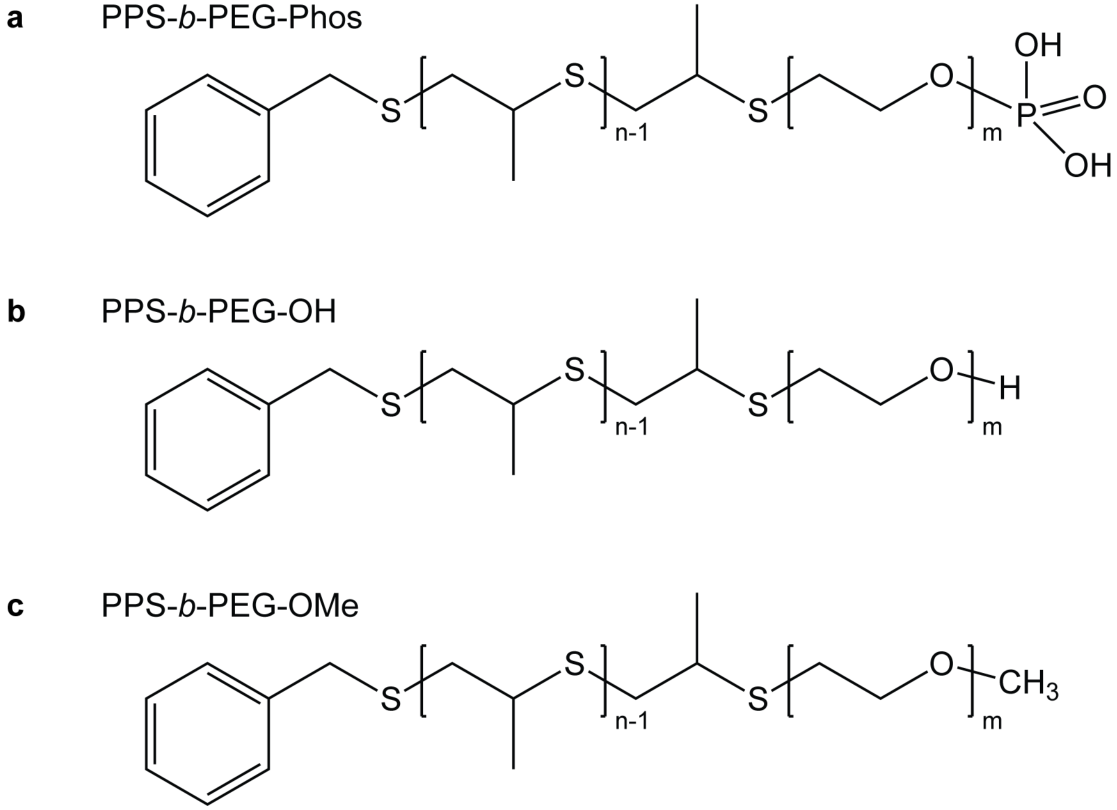


**Supplementary Fig. 1. | Self-assembling PEG-*b*-PPS diblock copolymers used in this study. a-c**, Chemical structure of the (**a**) Phos PS, (**b**) OH PS, and (**c**) MeO PS diblock copolymers.

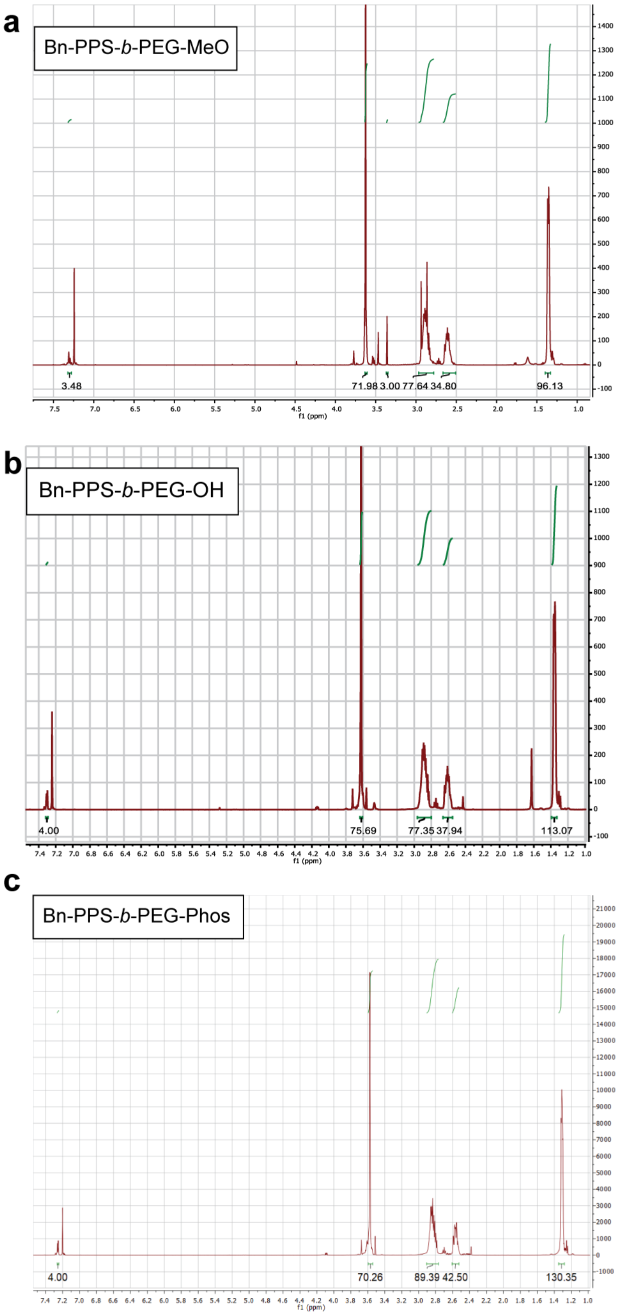


**Supplementary Fig. 2. | Sample $^1$H NMR Spectra of PEG-*b*-PPS polymers. a-c**, $^1$H NMR spectrum for MeO PS (**a**), OH PS (**b**), and Phos PS (**c**) polymers is displayed.

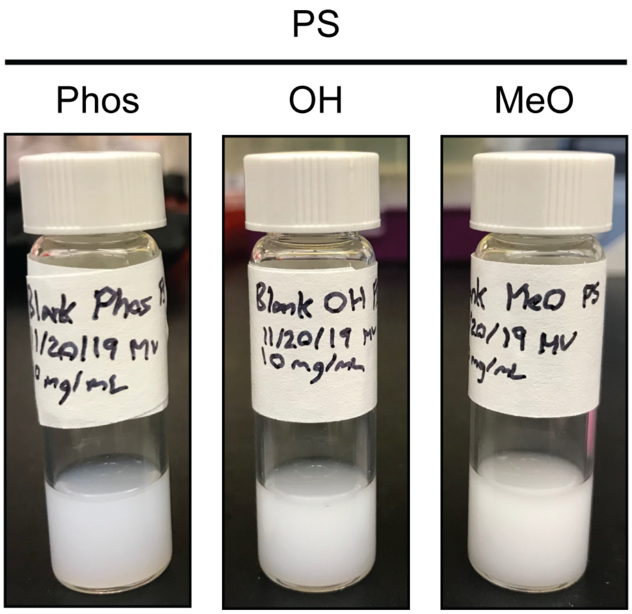


**Supplementary Fig. 3. | Images of representative PEG-*b*-PPS Phos, OH, and MeO polymersome formulations.** The polymer concentration is 10 mg/mL. The displayed formulations are not loaded with cargo (dyes, etc.).

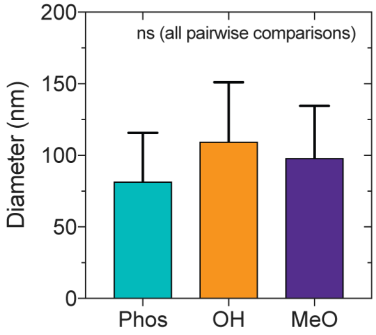


**Supplementary Fig. 4. | Comparison of PEG-*b*-PPS nanocarrier diameter.** The diameter of PEG-*b*-PPS polymersomes was measured by dynamic light scattering (DLS). Error bars represent s.d. (n = 3). Significance was determined by ANOVA with Tukey's multiple comparison test (5% significance level). All pairwise comparisons were found to be not significant (ns).

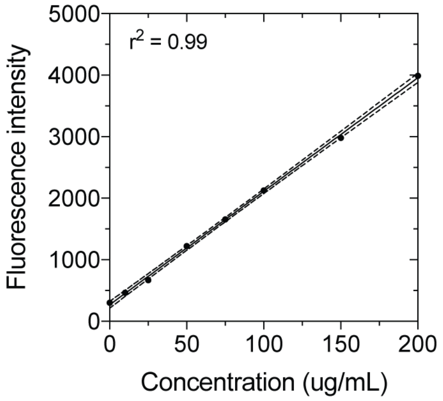


**Supplementary Fig. 5. | Dex$_{70\ kDa}$-TMR calibration.** Calibration curve prepared using a concentration series of 70 kDa dextran-tetramethylrhodamine (Dex$_{70\ kDa}$-TMR). Each standard was prepared in triplicate. The data were fit using a linear regression model, y = 18.4x + 270.2 ($r^2$ = 0.99). Error bars (s.e.m.) are not displayed due to being smaller than the height of the symbol. Bands represent the 95% confidence interval.

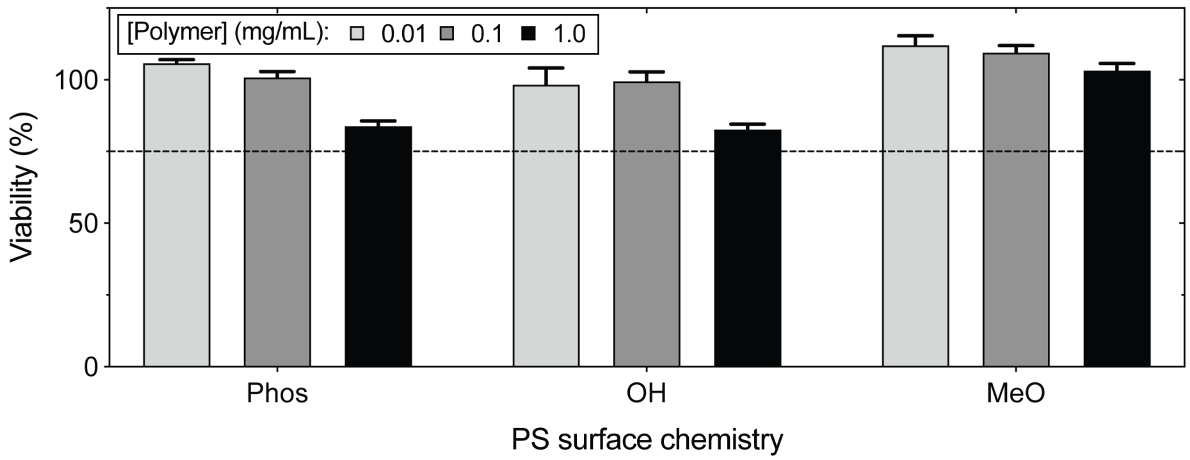


**Supplementary Fig. 6. | PEG-*b*-PPS polymersomes are non-toxic to RAW 264.7 macrophages *in vitro*.** RAW 264.7 macrophages cultured in CDMEM were exposed to Phos PS, OH PS, or MeO PS at a polymer concentration of 0.01 mg/mL, 0.1 mg/mL, or 1.0 mg/mL at 37 °C for 8 h. The MTT assay was used to assess viability (n = 5). The percentage viability was calculated with respect to a 1x PBS-treated control. The 75% viability line is indicated.

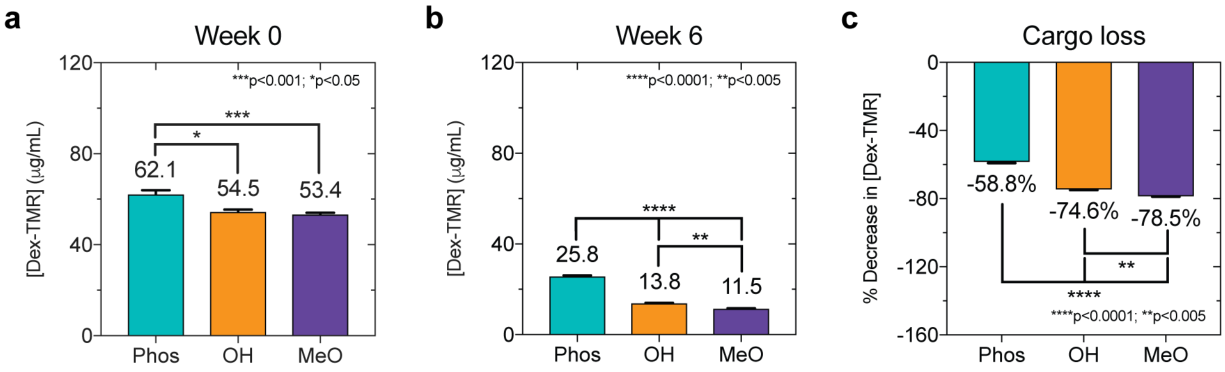


**Supplementary Fig. 7. | Hydrophilic cargo retention by PEG-*b*-PPS PS of different surface chemistry stored at 4 °C for 6 weeks. a**, **b**, Concentration (μg/mL) of $Dex_{70\ kDa}$-TMR at (**a**) 0 weeks determined after removing unencapsulated $Dex_{70\ kDa}$-TMR by size exclusion chromatography using a Sepharose 6B column, and (**b**) after storage at 4 °C for 6 weeks. At the 6-week timepoint, each PS formulation was sedimented by ultracentrifugation at 25,000 x g for 30 minutes under vacuum at 4 °C. A Beckman Coulter Optima Max-XP ultracentrifuge was used to isolate $Dex_{70\ kDa}$-TMR PS. Free (unencapsulated/lost) Dex-TMR was removed from the supernatant. The pellet (containing sedimented $Dex_{70\ kDa}$-TMR PS) was resuspended in 1x PBS to the input volume. **c**, The percentage decrease in $Dex_{70\ kDa}$-TMR concentration determined at the 6-week timepoint. In all plots, Phos, OH, and MeO correspond to the polymersome surface chemistry. Error bars represent s.e.m. Significance was determined by ANOVA with Tukey's multiple comparison test (5% significance level). *$p$<0.05, **$p$<0.005, ***$p$<0.001, ****$p$<0.0001.

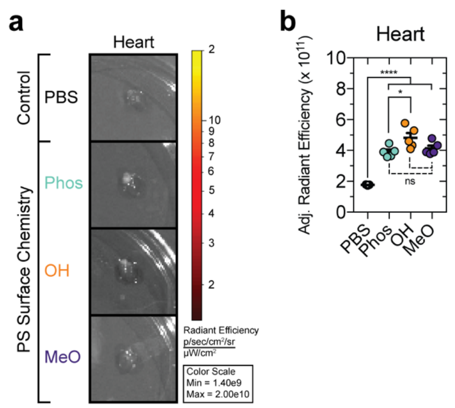


**Supplementary Fig. 8. | Detection of MeO PS, OH PS, and MeO PS in the heart 4 h after IV administration in. C57BL/6J mice. a**, Adjusted radiant efficiency measured in dissected heart tissue. **b**, Comparison of the adjusted radiant efficiency between treatment groups determined in heart tissue. Significance was determined using Tukey's multiple comparisons test with a 5% significance level. *****p*<0.0001, ****p*<0.0005, ***p*<0.01, **p*<0.05, ns = not significant.

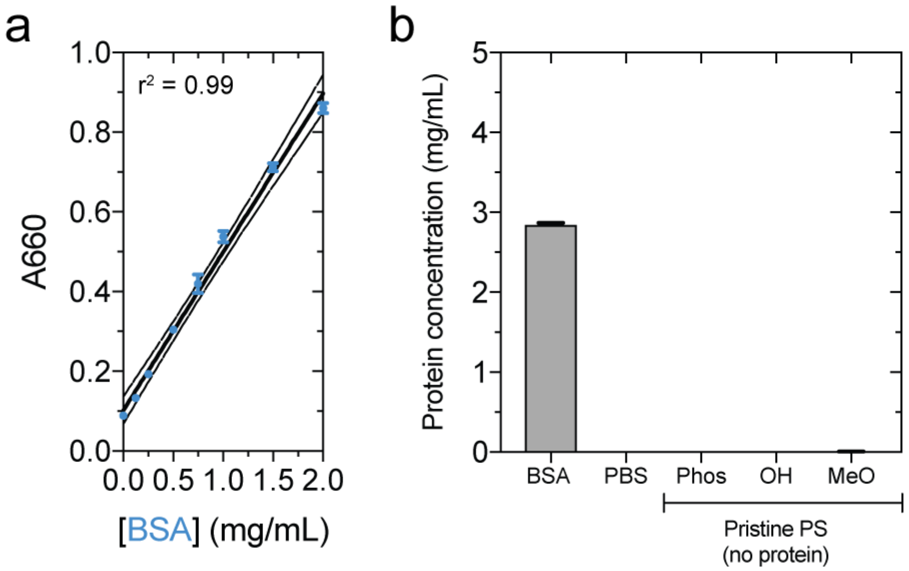


**Supplementary Fig. 9. | PEG-*b*-PPS polymers do not interfere with the Pierce A660 assay used to determine protein concentration.** **a**, Example BSA calibration curve used to determine the concentration of BSA adsorbed to PS-BSA complexes the Pierce A660 nm assay. Bands (dashed lines) represent the 95% confidence interval for the fit linear regression model (y=0.0003975x + 0.1019; $r^2$ = 0.99). **b**, Assessment of polymer interference on protein measurement. The concentration of a 0.2 μm-filtered BSA sample (prepared at 3.0 mg/mL) is shown together with a 1x PBS negative control (background). The signal (or lack thereof) from pristine Phos PS, OH PS, and MeO PS at a polymer concentration of 2.5 mg/mL is shown. The pristine condition refers to the absence of protein. Error bars represent s.e.m.

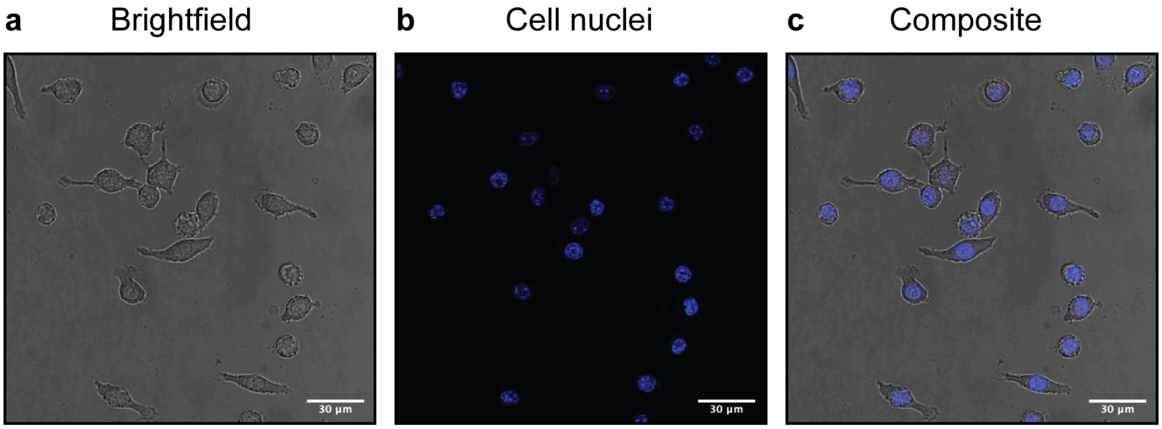


**Supplementary Fig. 10. | Live cell confocal images of RAW 264.7 macrophage morphology. a-c,** Representative (**a**) Brightfield, (**b**) stained cell nuclei (NucBlue), and (**c**) composite (brightfield/NucBlue overlay) confocal microscopy images are displayed for the RAW 264.7 macrophages used in this study. The objective magnification is 63X for all images. Scale bar = 30 μm.

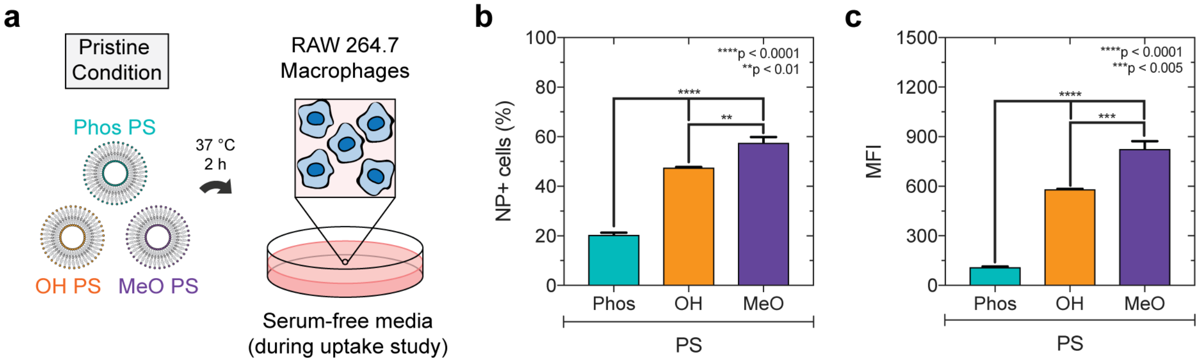


**Supplementary Fig. 11. | Statistical comparison of pristine polymersome uptake under serum-free conditions. a-c**, Baseline uptake of pristine PS and PS-BSA complexes by RAW 264.7 macrophages *in vitro*. **a**, Experiment illustration depicting macrophage treatment with pristine $Dex_{70\ kDa}$-TMR-loaded Phos PS, OH PS, and MeO PS nanocarriers for 2 h at 37 °C. The pristine condition refers to the absence of a pre-formed BSA corona. Cells were cultured in normal CDMEM media (DMEM +10% fetal bovine serum (FBS) +pen/strep) until 30 minutes prior to the uptake study. Cells were washed in PBS and were subsequently placed in DMEM containing pen/strep antibiotics but lacking FBS. This is a serum-free culture condition that allows the examination of the cellular uptake of each PS type in the absence of a protein corona. Cells were then treated with PS (0.5 mg/mL working polymer concentration) for 2 h at 37 °C. At that time, nanoparticles were removed via aspiration. The cells were subsequently washed, fixed, and stained prior to flow cytometric analysis (**b, c**). **b**, The percentage of nanoparticle-positive (NP+) cells determined using a false positive rate of <1.5%. The. false positive rate was defined using PBS-treated cells. **c**, The median fluorescence intensity (MFI) above background (untreated cells). Error bars represent s.e.m. Significance was determined by ANOVA with Tukey's multiple comparisons test. **$p$<0.01, ***$p$<0.005, ****$p$<0.0001.

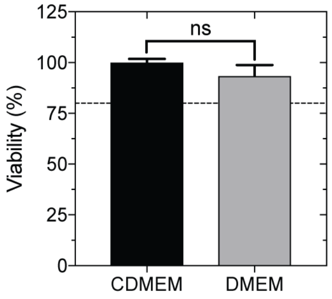


**Supplementary Fig. 12. | RAW 264.7 macrophage viability in serum-free media versus serum-supplemented media.** RAW 264.7 macrophages were incubated for 24 h at 37 °C in serum-free (DMEM) lacking FBS, or in complete DMEM (CDMEM) containing 10% FBS. Cell viability was assessed using the MTT assay, measuring the absorbance of 570 nm light (n = 5). The absorbance values of PBS-treated cells were normalized to the mean absorbance value of the CDMEM condition. Error bars represent the s.e.m. (n = 5). A t-test was performed to determine whether the mean values are significantly different between the two treatment conditions. ns = not significant (5% significance level).

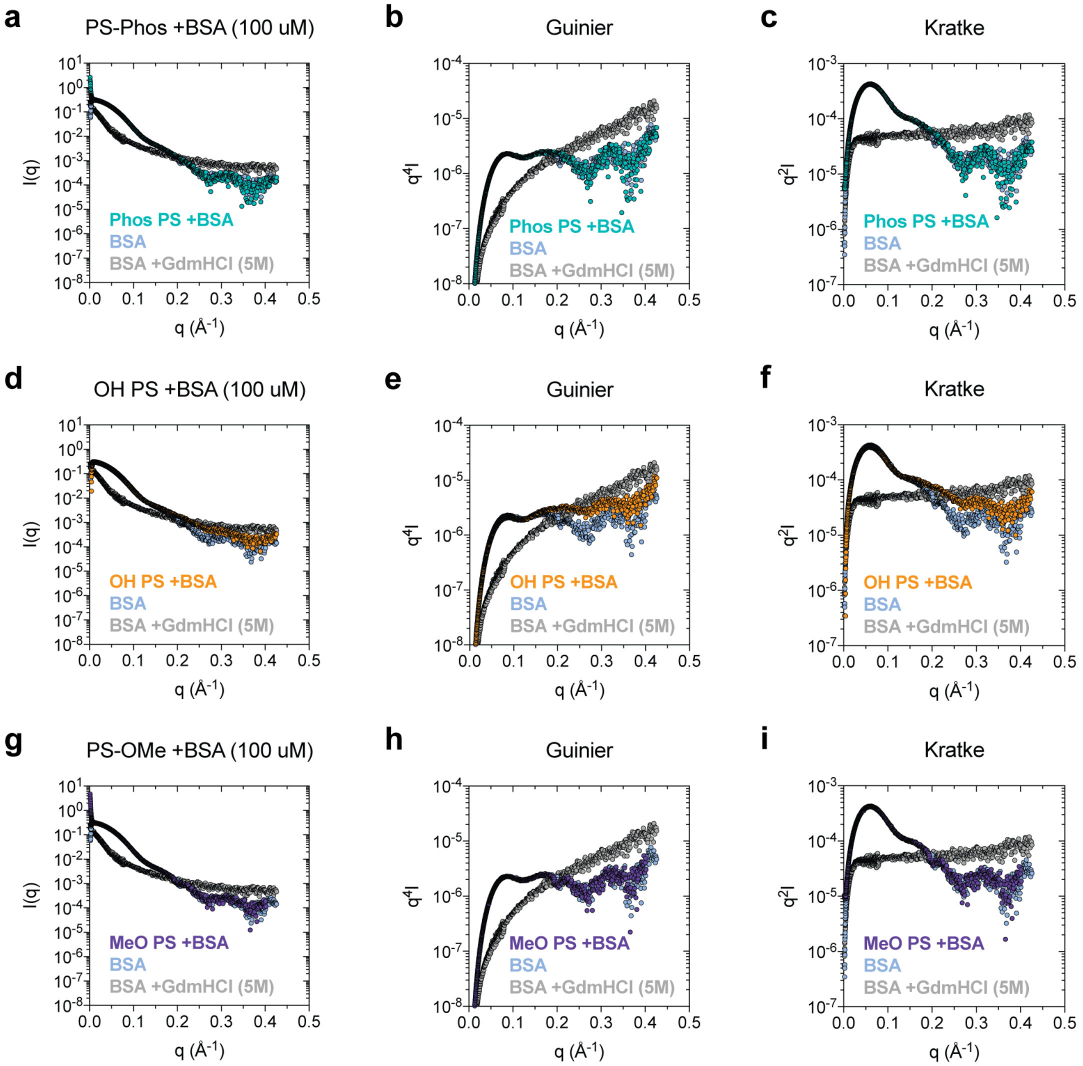


**Supplementary Fig. 13. | Small angle x-ray scattering of albumin adsorbed to Phos PS, OH PS, and MeO PS nanocarriers.** SAXS profiles, Guinier analyses, and Kratke analyses for BSA adsorbed to Phos PS (**a-c**), OH PS (**d-f**), and MeO PS (**g-i**). The average scattering profile obtained from 7 replicates is displayed in (**a, d, g**) for each sample. Background subtraction was performed by subtracting all non-protein contributions to the scattering (i.e. buffer, polymersomes, etc. depending on the sample). In all cases, folded BSA input (blue) and BSA denatured with 5M Gdm-HCl (gray) are displayed as references for the purpose of comparison.

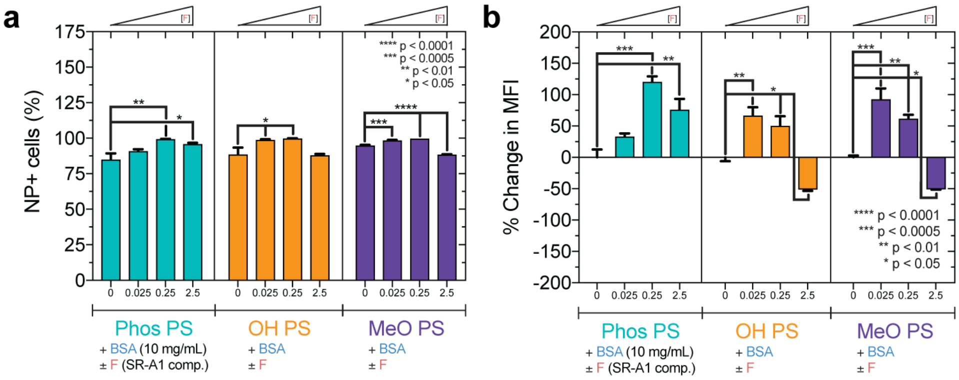


**Supplementary Fig. 14. | Concentration-dependence of fucoidan pre-treatment on macrophage uptake of PS-BSA complexes in serum-free media *in vitro*.** PS-BSA were formed in 10 mg/mL BSA. RAW 264.7 cells were treated with 0, 0.025, 0.25, or 2.5 mg/mL fucoidan (F) in serum-free DMEM. PS-BSA ([polymer] = 0.5 mg/mL) were administered after a 30 min incubation period. **a**, The percentage of NP+ cells. **b**, The percent change in median fluorescence intensity (MFI) comparing PS uptake in the absence versus presence of fucoidan. This percent change metric was calculated for each competitor concentration examined. Note that since this calculation employs the mean MFI determined in the absence of competitor, error in this mean value was accounted for by computing the percent difference between the individual MFI values measured for the specified PS-BSA (10 mg/mL) and its corresponding mean MFI. ANOVA with Dunnett's multiple comparisons test was performed to determine whether the uptake of PS-BSA complexes was significantly different in the presence of the specified concentration of Fucoidan (F) SR-A1 competitor. The p-values are $*p<0.05$, $**p<0.01$, $***p<0.0005$, and $****p<0.0001$. A false positive rate of 1.1% was used when computing the percentage of NP+ cells. The false positive rate was determined using PBS-treated cells (NP-). The reported MFI is the MFI above background, defined as the MFI measured for PBS-treated cells. Error bars represent the s.e.m (n = 3) unless indicated otherwise. In (**b**), the 2.5 mg/mL Fucoidan pre-treatment data is displayed from **Fig. 5** for the purpose of comparison.

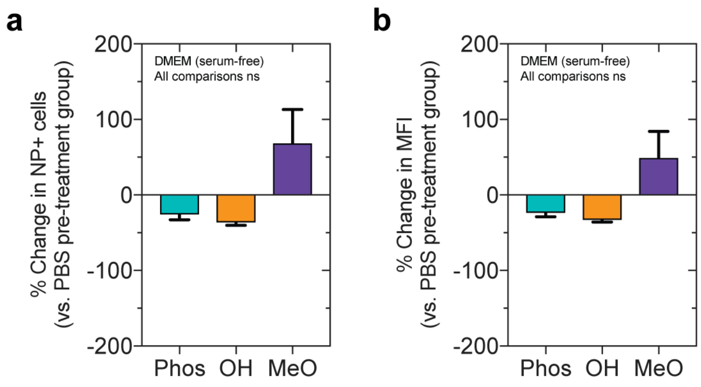


**Supplementary Fig. 15. | Macrophage uptake of Phos, OH, and MeO PS-BSA does not significantly change after pre-treating cells with polyinosinic acid in serum-free conditions.** Prior to assessing the uptake of Dex-TMR Phos PS, OH PS, and MeO PS with a pre-adsorbed BSA corona, RAW 264.7 macrophages were pre-treated with 2.5 mg/mL polyinosinic acid prepared in DMEM (serum-free). The percent change in (**a**) the percentage of NP+ cells (false positive rate < 1.5%), and (**b**) the median fluorescence intensity (MFI) above background (PBS-treated cells) is displayed. Statistical significance was determined using ANOVA with Tukey's multiple comparisons test (5% significance level). All pairwise comparisons were not significant (ns).

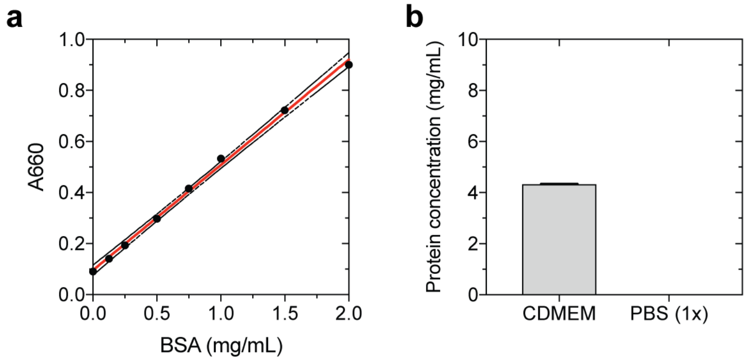


**Supplementary Fig. 16. | Protein concentration in complete DMEM used in serum containing conditions. a**, Pierce A660 assay calibration against a BSA concentration series. A linear regression model was fit (y = 0.4124x + 0.09569) with an $r^2$ value of 0.99. The fit line is shown in red together with the 95% confidence interval (dashed black lines). The mean of the measured values is plotted. BSA standards were prepared in triplicate. Error bars represent the s.e.m. (n = 3). If error bars are not shown, this indicates the error is so small that the error bars would not be visible beyond the symbol plotted for the mean value. **b**, The protein concentration of complete DMEM (CDMEM) used in this study. CDMEM is supplemented with 10% fetal bovine serum (FBS). The protein concentration in this media was determined to be 4.33 ± 0.01 mg/mL. PBS is included as a negative control.

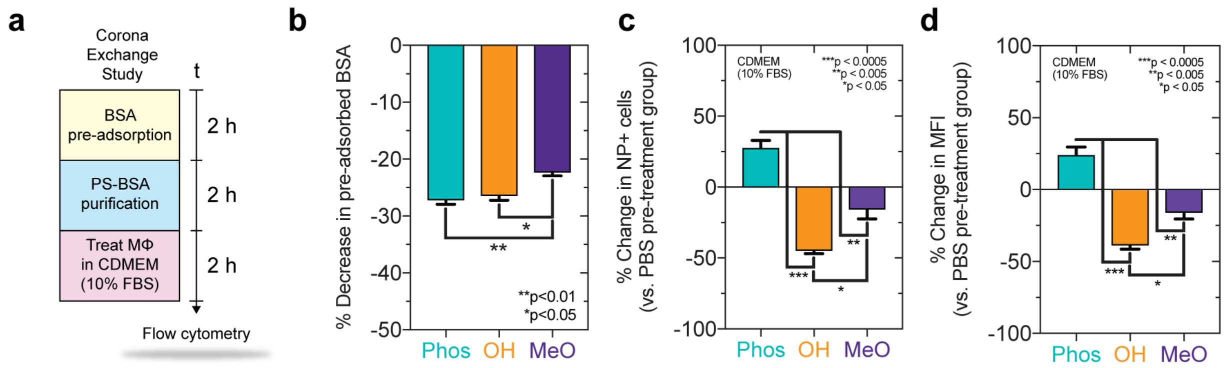


**Supplementary Fig. 17. | Macrophage uptake of PS-protein complexes after exchanging pre-adsorbed BSA with serum proteins. a**, Uptake study illustration. BSA was pre-adsorbed to PS. PS-BSA complexes were purified by ultracentrifugation and were subsequently incubated with RAW 264.7 cells cultured in CDMEM containing 10% FBS. **b**, The percent decrease in the amount of pre-adsorbed BSA after incubation in CDMEM (containing 10% FBS) at 37 °C for 2 h. For the purpose of quantifying adsorbed protein exchange in this particular experiment, FITC-BSA was used as a proxy for unlabeled BSA to quantify its loss after adsorption. Separate PS samples were prepared in parallel with FITC-BSA (for exchange quantification) and unlabeled native BSA (for samples used in the actual uptake studies). ANOVA with Tukey's multiple comparisons test was used to determine statistical significance. *$p<0.05$, **$p<0.01$. **c**-**d**, Flow cytometric analysis of the percent change in (**c**) the percentage of NP+ cells and (**d**) the MFI. The percent change was calculated between the +/- Fucoidan (2.5 mg/mL) groups. Significance was determined using ANOVA with Tukey's multiple comparisons test (5% significance level). *$p<0.05$, **$p<0.005$, ***$p<0.0005$. Error bars represent the s.e.m. for all cases (n = 3).
